# A new twist of rubredoxin function in *M. tuberculosis*

**DOI:** 10.1101/2020.10.27.356691

**Authors:** Tatsiana Sushko, Anton Kavaleuski, Irina Grabovec, Anna Kavaleuskaya, Daniil Vakhrameev, Sergei Bukhdruker, Egor Marin, Alexey Kuzikov, Rami Masamrekh, Larisa V. Sigolaeva, Victoria Shumyantseva, Kouhei Tsumoto, Valentin Borshchevskiy, Andrei Gilep, Natallia Strushkevich

**Affiliations:** The Institute of Medical Science, the University of Tokyo, Tokyo, Japan; Institute of Bioorganic Chemistry, National Academy of Sciences of Belarus, Minsk, Belarus; Research Center for Molecular Mechanisms of Aging and Age-Related Diseases, Moscow, Institute of Physics and Technology (MIPT), Dolgoprudny, Russia; Institute of Biomedical Chemistry, Moscow, Russia; Pirogov Russian National Research Medical University, Moscow, Russia; Department of Chemistry, M.V. Lomonosov Moscow State University, 119991 Moscow, Russia; Department of Bioengineering, School of Engineering, the University of Tokyo, Tokyo, Japan; Skolkovo Institute of Science and Technology, Moscow, Russia

**Keywords:** rubredoxin, *M. tuberculosis*, electron transfer, cytochrome P450, CYP, redox partner

## Abstract

Electron transfer mediated by metalloproteins drives many biological processes. Rubredoxins are ubiquitous iron-containing electron carriers that play important roles in bacterial adaptation to changing environmental conditions. In *Mycobacterium tuberculosis*, oxidative and acidic stresses as well as iron starvation induce rubredoxin expression. However, their functions during *M. tuberculosis* infection is unknown. In the present work, we show that rubredoxin B (RubB) supports catalytic activity of mycobacterial cytochrome P450s, CYP124, CYP125, and CYP142, which are important for bacterial viability and pathogenicity. We solved the crystal structure of RubB and characterized the interaction between RubB and CYPs using site-directed mutagenesis. Mutations that neutralized single charge on the surface of RubB did not dramatically decrease activity of studied CYPs, and isothermal calorimetry (ITC) experiments indicated that interactions are transient and not highly specific. Our findings suggest that a switch from ferredoxins to rubredoxins support CYP activity in *M. tuberculosis*-infected macrophages. Our electrochemical experiments suggest potential applications of RubB in biotechnology.

## Introduction

Tuberculosis was the first infectious disease to be declared a global health emergency by the World Health Organization; tuberculosis causes more than 1,7 million deaths every year. Multiple mechanisms allow *Mycobacterium tuberculosis* (Mtb), the bacterium that causes tuberculosis, to survive within macrophages. These mechanisms include production of the antioxidant molecule mycothiol [1], activation of the catalase-peroxidase katG and superoxide dismutases sodA and sodC [2], and synthesis of truncated hemoglobins [3]. In phagosomes, Mtb resides in an acidic environment, thus requiring mechanisms to sustain growth under oxidative and acidic stresses. Moreover, during granuloma formation Mtb experiences drastic iron deprivation that influences the intensity of iron-sulfur clusters assembly in redox proteins [4]. The presence of a variety of genes that encode [3Fe-4S] and [4Fe-4S] ferredoxins and rubredoxins identified in Mtb suggests the important role of iron-containing proteins in maintaining redox homeostasis. We hypothesized that rubredoxins, which have been suggested to be part of an evolutionary chain between ferredoxins and flavodoxins [5], might be important in reactions catalyzed by cytochrome P450 (CYP) proteins.

Rubredoxins are small (∼6 kDa) iron–sulfur proteins that are crucial for oxidative stress responses. They rapidly transfer metabolic reducing equivalents to oxygen or reactive oxygen species and act as electron carriers in many biochemical pathways. The Mtb genome contains a highly conserved operon comprised of two tandemly arranged rubredoxin-encoding genes, *RubA* (Rv3251c) and *RubB* (Rv3250c), and the gene *alkB* (Rv3252c), which encodes an alkane hydroxylase. The gene products are probably involved in alkane and fatty acid metabolism. Our bioinformatic analysis of rubredoxins across different phylogenetic groups (Fig.S1, S2) showed that *RubA* is conserved in *Actinobacteria* and is specific for this family, whereas *RubB* is observed in gamma- and beta-proteobacteria. Large and distinctive rubredoxin families are found in *Bacteroidetes* and *Firmicutes* phyla; these genes group with genes encoding primitive archaeal rubredoxins, and plant-specific rubredoxins, demonstrating evolutionary conservation of rubredoxin scaffold.

Expression of both Mtb rubredoxins is induced under iron starvation [4] and during iron chelator 2,2’-bipyridyl application [6]. Oxidative stress induced by the alkaloid ascidemin or nitrosative stress caused by diethylenetriamine/NO or a mixture of S-nitrosoglutathione and potassium cyanide strongly induce expression of rubredoxins in Mtb culture. Finally, rubredoxins are also induced by culture of Mtb in acidic pH [6, 7]. Notably, rubredoxin induction is often accompanied by inhibition of ferredoxin expression.

Induction of rubredoxin expression during oxidative and acidic stresses as well as under iron starvation might be due to the physical-chemical properties of rubredoxins. Rubredoxins need only one iron ion, which is beneficial during residence within macrophages in which microelements are deficient. Further, rubredoxins are usually of lower molecular mass than ferredoxins and do not require scaffold proteins, thus requiring few biosynthetic resources than ferredoxins. Moreover, rubredoxin activity is not inhibited by oxidative, acidic and temperature stresses, which may be crucial in the face of a host immune response [8, 9]. This evidence suggests that rubredoxins more advantageous redox partners for CYPs than ferredoxins.

To test this idea, we purified RubB protein and measured enzymatic activity of three Mtb CYPs in the reconstituted system, containing RubB and different reductases as redox-partners. Our results demonstrate that RubB supports CYP-dependent catalysis. We also performed biophysical characterization of RubB including crystal structure determination. Site-directed mutagenesis was used for mapping of the protein-protein interactions within the RubB - CYP complex. Based on data obtained using purified RubB mutants, we suggest that the RubB interaction with CYPs is transient and not highly specific, as point charge neutralization on the surface of RubB does not dramatically affect CYP activity. Overall, our results provide new insights into electron transfer within different classes of redox proteins in Mtb and suggest that these interactions might be important switch mechanisms during different stages of Mtb infection.

## Results

### Cloning, expression and purification of RubB

*RubB* was cloned into the pET11a vector and overexpressed in *Escherichia coli*. According to a previous report, both iron and zinc-substituted rubredoxins are produced during heterologous expression in *E.coli* [10]. As binding of zinc and iron to RubB might be a competitive process [11], we provided extra iron during culture to obtain primarily the iron-containing form. RubB was purified to a homogenous state using sequential ion-exchange and size-exclusion chromatography. Based on size-exclusion chromatography, RubB was predominantly in the monomeric form and had spectral properties typical of oxidized Fe-S rubredoxins with maxima at 280, 380, and 490 nm (Fig. 1).

**Fig. 1.**
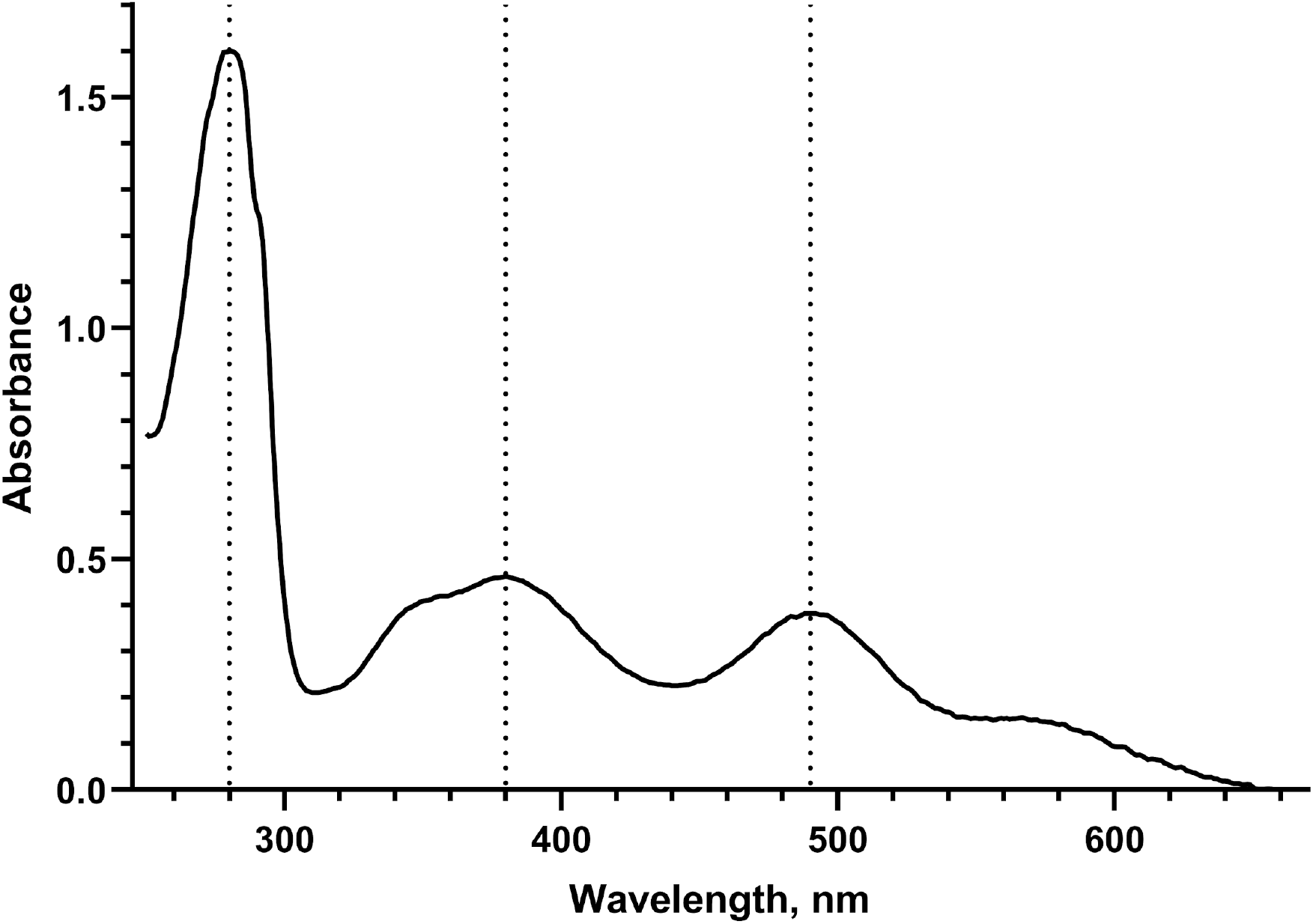
UV-Vis absorbance spectrum of RubB.

### Thermal stability of RubB

Some rubredoxins are hyper thermostable [12]. Previous circular dichroism (CD) studies of zinc-substituted Mtb RubB revealed that RubB is stable to 80°C; however, 26% of residues of that protein are in the unstructured N-terminal tag, and its unfolding temperature was not determined [10]. Our analysis of thermal stability of purified RubB from Mtb using differential scanning calorimetry (DSC) revealed no peaks indicative of thermal unfolding in the temperature range 10-110 °C over the pH range from 6 to 7.4 (Table 1). A constant increase in heat capacity due to hydration of residues was observed over this temperature range. Thermal stability at acidic pH, conditions under which protein side chain are protonated, was more stable at pH 5 (melting temperature (T_m_) of 87.2°C) than at higher or low pH (Table 1). The CD-spectra of RubB measured at pH values from 5.0 to 7.4 were not significantly different. Thus, RubB has a native fold and high thermal stability in the pH range (pH 4.5-6.2) of the phagosomes of macrophages where Mtb resides [9]. Our data are in good correlation with heat-induced denaturation of rubredoxins from other sources [13].

**Table 1.** Influence of pH on thermal stability of RubB estimated using DSC

| pH | 2.0 | 3.0 | 4.0 | 5.0 | 6.0 | 7.4 |
| --- | --- | --- | --- | --- | --- | --- |
| T<sub>m</sub>, °C | 57.4±0.6 | 60.4±0.8 | 70.1±0.9 | 87.2±0.7 | nd | nd |

nd – T<sub>m</sub> could not be determined

### Electrochemistry

Rubredoxins are involved in the electron transfer processes. These proteins cycle between ferric and ferrous states [14]. Rubredoxins of different organisms have different redox potentials ranging from −100 to +50 mV. Depending on redox potential, rubredoxins are divided into low potential and high potential groups. To study the redox potential of Mtb RubB, we evaluated the direct electron transfer process on an electrode surface as non-catalytic cycling between oxidized ferric and reduced ferrous states. A solution of RubB in the supporting electrolyte gave no cyclic voltammetric response at bare screen-printed electrode (SPE). However, modification of the SPE with multiwalled carbon nanotubes (MWCNTs) led to well-defined cyclic voltammograms (Fig.2A).

**Fig 2.**
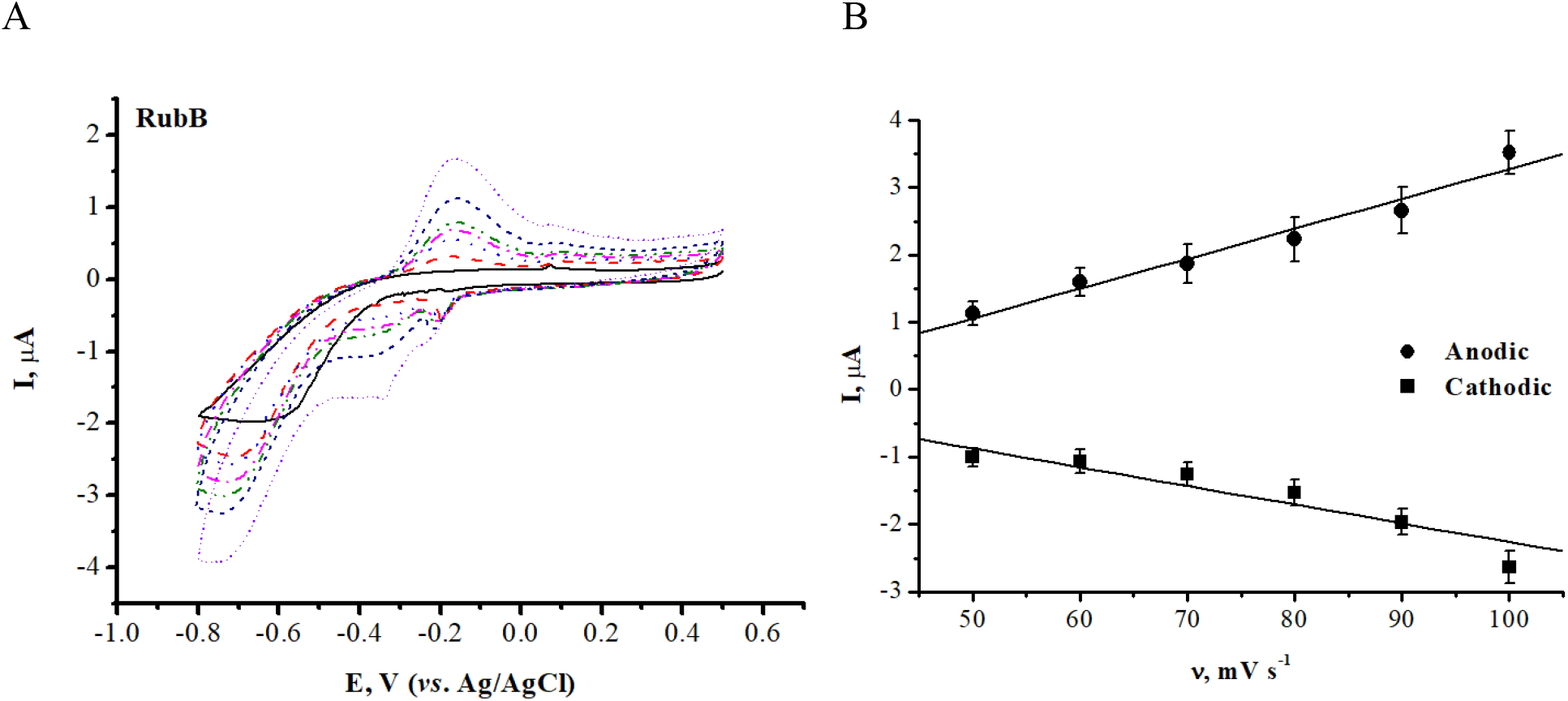
(A) Direct non-catalytic cyclic voltammograms of MWCNT-modified SPE without (solid lines) and with RubB (dashed lines) in 0.1 M potassium phosphate buffer (pH 7.4) containing 0.05 M NaCl from 50 to 100 mV/s. (B) Dependence of cathodic and anodic current on scan rate.

Direct electrochemistry of RubB was achieved by noncovalent immobilization on an MWCNT-modified SPE. Two peaks of the iron-sulfur cluster characterizes reduction and oxidation of RubB. Analysis of voltammograms revealed behavior consistent with quasi-reversibility; indeed, with increased scan rate there was an increase in separation between anodic and cathodic peaks (Fig. 2B). The observed linear dependence of anodic and cathodic peak current on the scan rate indicated that the electron transfer from and to the electrode is a reversible and surface-controlled process. The calculated value of the electrochemical potential of RubB suggest that electron transfer from Mtb reductase FprA is possible (Table 2). Our results are in accordance with previously reported data on electrochemistry of rubredoxins obtained using gold disk or carbon electrodes modified by means of neomycin and Mg^2+^ ions [15].

**Table 2.**
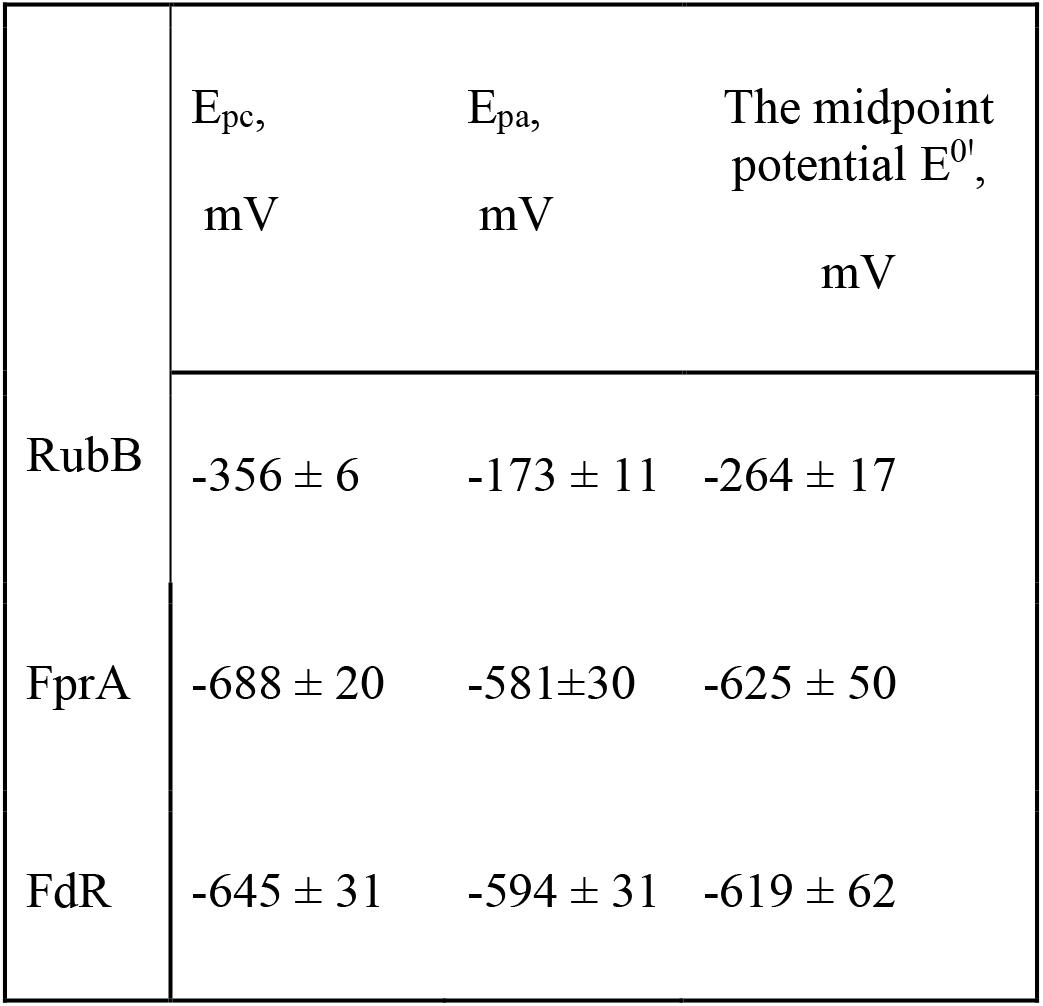
Electrochemical parameters of RubB, FprA, and FdR. Analyses were performed on an SPE modified with MWCNT in 0.1 M potassium phosphate buffer (pH 7.4), containing 0.05 M NaCl. Epc is the signal at the cathodic peak, Epa is the signal at the anodic peak, and E^0’^ is the midpoint potential.

### Reconstitution of CYP activity

Mtb CYPs, like most bacterial CYPs, belong to the class I electron-transfer system and are driven by ferredoxin and ferredoxin reductases that transfer the electrons from the NAD(P)H to the CYP heme cofactor. Recently, it was shown that activity of several Mtb CYPs could be supported by specific cognate redox partners *in vitro* [16], but redox partners are still unknown for most Mtb CYPs.

It was shown that RubB is co-expressed with CYP125 in a macrophage infection model [17] and with CYP142 during the transition to dormancy [18]; these data link RubB to cholesterol metabolism. To evaluate the ability of RubB to support enzymatic activity of CYPs involved in cholesterol metabolism, we selected CYP124, CYP125, and CYP142 as these are enzymes with well-established catalytic profiles and crystal structures are available [19]. The enzymatic reaction was performed a reconstituted system, containing reductase, RubB, and CYP. Analysis of high performance liquid chromatography (HPLC) profiles clearly demonstrate the formation of the product for all tested CYPs, indicating that recombinant RubB transferred electrons from the reductase to CYP. The amounts of products formed in the presence of RubB were comparable to that produced when the most efficient system with spinach ferredoxin was used [20]. Products were formed with significantly different efficiencies when reductases from various sources were used (Table 3), demonstrating the selectivity of RubB. The most efficient redox-partners for RubB in CYP124-catalyzed reaction were reductases Arh1 from *Saccharomyces cerevisiae* and PETH from *Spinacia oleracea,* whereas cognate reductases NADH-ferredoxin reductase FdR and NADPH-ferredoxin reductase FprA resulted in considerably less product. In contrast, for CYP125 the preferred reductase was FprA. Thus, Rub efficiently mediates the hydroxylation reactions of cholesterol-metabolizing CYPs and the product conversion rates were modulated by the reductase component of the system.

**Table 3.** Enzymatic activity of CYPs in the reconstituted system containing RubB and reductases. If RubB alone or reductase alone were added to CYP and NAD(P)H, catalytic activity was not detected. CYP124 catalytic activity was measured by oxidation of 7-ketocholesterol. CYP125 and CYP142 activities were measured by oxidation of cholestenone.

| Reconstituted system | Enzymatic activity, min<sup>-1</sup> |  |  |
| --- | --- | --- | --- |
|  | CYP124 | CYP125 | CYP142 |
| Arh1 (from *Saccharomyces cerevisiae*), RubB | 4.23±0.06 | 0.36±0.04 | 1.08±0.01 |
| FprA (from *Mtb*), RubB | 2.35 ± 0.16 | 0.52 ±0.02 | 0.98 ±0.20 |
| FdR (from *Mtb*), RubB | 2.49±0.03 | 0.09±0.03 | 0.48±0.11 |
| PETH (from *Spinacia oleracea*), RubB | 5.25± 0.42 | 0.56±0.30 | 1.01±0.09 |

### Crystal structure of RubB

The crystal structure of RubB was determined with anisotropic resolution (1.2 × 1.3 × 1.6 Å). The crystals belong to the P1 space group with eight molecules in the asymmetric unit. All amino acids were built except C-terminal residues A58-S60 of three monomers (in chains B, D and H). Protein molecules in the asymmetric unit did not have significant structural differences with the Cα root mean square deviation (RMSD) not exceeding 0.7 Å (Fig. 3).

**Fig 3.**
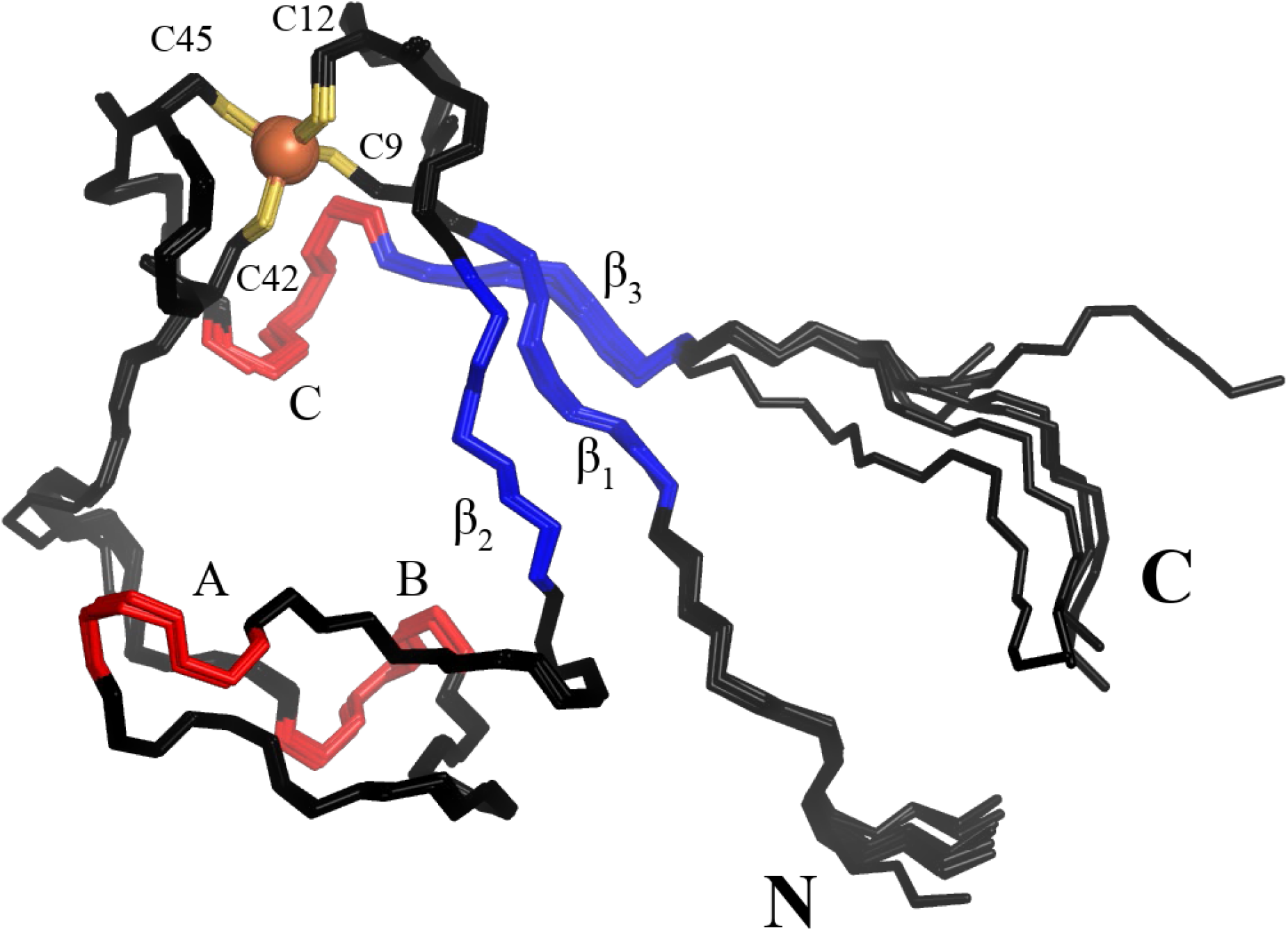
Structures of the eight monomers of RubB in the asymmetric unit of the crystal structure. Aligned structures of Cα bonds are shown as sticks. The central β-sheet is colored blue, the 310-helices (denoted A, B and C) are colored red. In the FeS4 cluster, the iron ion is shown as an orange sphere, and sulfurs of the coordinating cysteines are depicted in yellow. The differences between monomers in the asymmetric unit are determined to a great extent by M1 at the N-terminus and E56-S60 at the C-terminus, for which average pairwise RMSD is 4.95 Å; for other residues pairwise RMSDs do not exceed 0.44 Å. This result is consistent with previously solved NMR structure, where residues D3-E56 had 0.39 Å RMSD for backbone atoms.

RubB has a rubredoxin fold similar to the fold previously characterized [21]. In particular, the RubB fold consists of one antiparallel β-sheet with three strands, which are connected by two loops. Each loop contains a conserved cysteine motif (CXXCG). Cysteines from the loops (Cys9, Cys12, Cys42, and Cys45) surround an iron ion in tetrahedral geometry. This FeS_4_ cluster is the active site of the protein. Average Fe-S bond lengths are similar to the typical value of 2.3 Å (2.33 Å, 2.31 Å, 2.33 Å, 2.30 Å with 0.03 Å standard deviation between 8 protein copies), indicating the iron is in the oxidized state [21].

Previously, a zinc-substituted RubB structure was solved by NMR (PDB ID: 2KN9). Pairwise Cα RMSD values between molecules in our X-ray structure and the NMR-based model do not exceed 1.1 Å. In the NMR model, residues at C and N termini show backbone variability, while the rest of the structure remain almost unperturbed. There are two major differences in the active sites between the two structures. The first is a rotameric state of D44. The second is that the bond lengths in FeS4 cluster are significantly longer in the NMR model (2.41 Å, 2.41 Å, 2.42 Å, 2.41 Å). The latter might result from the zinc substitution, as was previously observed for other rubredoxins by different techniques [22, 23].

### Role of zinc in structure and function of RubB

In the crystal structure, we observed a number of intermolecular and intramolecular contacts with zinc ions (Fig. S4) that may be a consequence of Glu and Asp on the surface. Coordination spheres in these contacts are formed by various protein and solvent groups, including carboxyl groups of Glu, Asp, acetate ions, and the C-terminal Ser, carbonyl and amino groups of N-terminal Met, and glycerol.

To determine whether zinc ions influence protein activity or stability we performed CD, ITC, and DSC studies. The CD spectrum of mycobacterial RubB is characterized by a double minimum at 204 and 227 nm (Fig. S3). The spectrum indicates that protein has not only canonical β-strands and helices, but also unstructured regions; this is in correspondence with the crystal structure. Addition of zinc did not induce changes in the CD spectrum. Zinc binding was further studied using ITC. The following parameters were estimated: K_d_ = 12.0±0.7μM, ΔH = 30.4±0.9 kcal/mol, -TΔS = −36.9±1 kcal/mol, and ΔG = −6.5±0.3 kcal/mol (Fig. S5). The obtained data demonstrated that zinc binds to RubB with high affinity and that binding is entropy-driven and thermodynamically favorable.

Next, we measured the activity of CYP124 in the presence of increasing zinc concentrations with another doubly charged metal ion (Ni^2+^) as reference (Table 4). A dramatic decrease of activity was observed as the zinc concentration was increased with almost no activity observed at the 100 μM Zn^2+^ in a reaction mixture with CYP124, RubB, and Arh1 reductase. In comparison, a 10 mM Ni^2+^ concentration decreased activity only 2 fold compared to activity in the absence of added ion. These experiments do not allow us to conclude which components of the redox system are affected by the presence of zinc. The interaction between redox-partners and/or direct intersection with the catalytic cycle of CYP may be inhibited by the specific metal “poisoning” effect.

**Table 4.** Effect of Zn^2+^on catalytic activity of CYP124.

|  | Zn<sup>2+</sup>, mM |  |  |  |  | Ni<sup>2+</sup>, mM |
| --- | --- | --- | --- | --- | --- | --- |
|  | 0 | 0.1 | 0.5 | 1 | 10 | 10 |
| Enzymatic activity, % of substrate conversion | 50.1 | 0.3 | 0.3 | 0.4 | 0 | 25.4 |

Since zinc ions participate in crystal contacts (Fig. S4), we hypothesized that RubB might become disfunction because of oligomerization in the presence of high concentrations of Zn^2+^. We performed size-exclusion chromatography-multiple angle light scattering analysis of RubB in the presence and absence of Zn^2+^. In both the presence and absence of Zn^2+^, RubB a homogeneous peak was observed with a mass of 6.6±0.3 kDa, corresponding to monomer. Thus, RubB does not appear to form dimers or oligomers in the presence of zinc. We further analyzed thermal stability of RubB in presence of increasing Zn^2+^ concentrations (Table 5). Only at high concentrations (10 mM Zn^2+^) were two unfolding events detected. The observed decrease in stability might be explained by the partial iron substitution of zinc for iron that occurs during heating.

**Table 5.** Influence of zinc on thermal stability of RubB.

| Zn<sup>2+</sup> concentration,<br>mM | T<sub>m1</sub>,<br>°C | T<sub>m2</sub>,<br>°C |
| --- | --- | --- |
| 1 | 54.7 | 87.4 |
| 5 | 62.4 | 93.4 |
| 10 | 65.7 | 98.0 |

### Isothermal calorimetry analysis of the interaction between RubB and CYP124

To characterize complex formation between CYP124 and RubB from a thermodynamic point of view, entropy, enthalpy, and Gibbs energy values were estimated by the ITC. RubB binds to CYP124 with dissociation constant of 12.8 μM. Complex formation is thermodynamically favorable and entropy and enthalpy driven (Table 6). To probe if the substrate binding induces conformational changes in CYP124 that influence binding of RubB, we performed ITC measurements in the presence of the CYP124 substrate cholestenone. Addition of cholestenone decreased affinity of CYP124 for RubB (Table 6), indicating that RubB preferentially binds to the ligand-free CYP124. The same effect was observed upon the interaction between electron transfer proteins P450cam and Pdx in *Pseudomonas putida* [24]. A decrease in affinity of CYP124 for RubB was also observed when the ionic strength of a solution was increased, indicating that electrostatic interactions stabilize the complex.

**Table 6.** Thermodynamic parameters for RubB binding to CYP124.

|  | K<sub>d</sub>,<br>μM | ΔH,<br>kcal/mol | -TΔS,<br>kcal/mol | ΔG,<br>kcal/mol |
| --- | --- | --- | --- | --- |
| CYP124 | 12.8±0.7 | -1.8±0.2 | -4.5±0.1 | -6.3±0.1 |
| CYP124 plus cholestenone | 20.3±1.2 | -1.7±0.1 | -4.5±0.1 | -6.2±0.1 |
| CYP124 plus 0.15 M NaCl | 61.8±3.5 | -29±3.0 | 23.5±1.9 | -5.5±0.2 |

### Probing protein-protein interactions by site-directed mutagenesis

In order to identify specific amino acids of RubB important for the activity, we first performed multiple sequence alignment (Fig. S6). The residues Q11, W22, S41, and D44 were outliers. It was previously proposed that L41 in *Clostridium pasteurianum* rubredoxin, which corresponds to D44 in RubB, acts as a charge-dependent gate to control solvent access to the FeS cluster and facilitate electron transfer [25]. Moreover, D44 conformations are different in our crystal structure than in the reported NMR structure (Fig. S7), suggestive of the functional relevance of this residue. We prepared RubB mutants in which residues were changed to those in other rubredoxins (Q11V, W22D, S41V, and D44A) and in which charge-neutralizing changes were made (E24A, D25A, D34A, D35A, D38A, and D39A). None of these mutants had characteristics different from the wild type RubB during expression and purification.

To assess whether mutations affected the structure of RubB, we performed CD. Comparison of spectra of wild type and mutant RubB indicated that mutations did not induce significant structural perturbations. We next examined effects of the mutations on thermal stability of RubB (Fig. S8). In all cases, DSC data fit a two-state model within the expected error of data collection. E24A and D25A mutations did not have an effect on RubB thermal stability. D44A, S41V, Q11V mutations increased the thermal transition temperature (+5.5 °C, +1.7 °C, +2.4 °C relative to wild type, respectively). The profound effect of the D44A mutation on thermal stability indicates that charge neutralization in this position increases overall stability of RubB (Table 7).

**Table 7.** Thermodynamic parameters for wild type and mutant RubB. Thermal stability was determined by DSC in 20 mM citrate buffer, pH 5.0.

| RubB | T<sub>m</sub>,<br>°C | ΔH,<br>kcal/mol |
| --- | --- | --- |
| Wild type | 87.2 | 119.8 |
| E24A | 87.9 | 117.8 |
| D44A | 92.7 | 120.1 |
| D25A | 86.9 | 118.2 |
| S41V | 88.9 | 117.5 |
| Q11V | 89.6 | 122.6 |

We next measured catalytic activity of CYP124 in the presence of RubB mutants in reconstituted system containing Arh1, FprA, and FdR (Fig. 4). A 2-fold difference between the wild type and mutant was considered a significant effect. When the Mtb reductase FdR was used, no significant decrease in enzymatic activity of CYP124 was detected for any RubB mutant relative to that in the presence of wild type RubB. In the case of Mtb reductase FprA and the heterologous redox protein Arh1, the D44A mutation resulted in a decrease of CYP124 activity by 2.2-fold, whereas other mutations had no effect. It is possible that D44 of RubB interacts with the reductase. To test this, we superimposed our crystal structure of RubB with that of the complex of rubredoxin and rubredoxin reductase from *Pseudomonas aeruginosa* [PDB ID: 2V3B]. The residues E24 and D44 of RubB correspond to E21 and D41 of rubredoxin from *P. aeruginosa.* These two residues form salt bridges with the reductase [26]. The rubredoxin-rubredoxin reductase complex has high charge complementarity yet is transient. This comparison suggests that effects of corresponding mutation at D44 of RubB could be explained by the impaired interactions with solvent exposed residues of FprA or Arh1 reductases.

**Fig 4.**
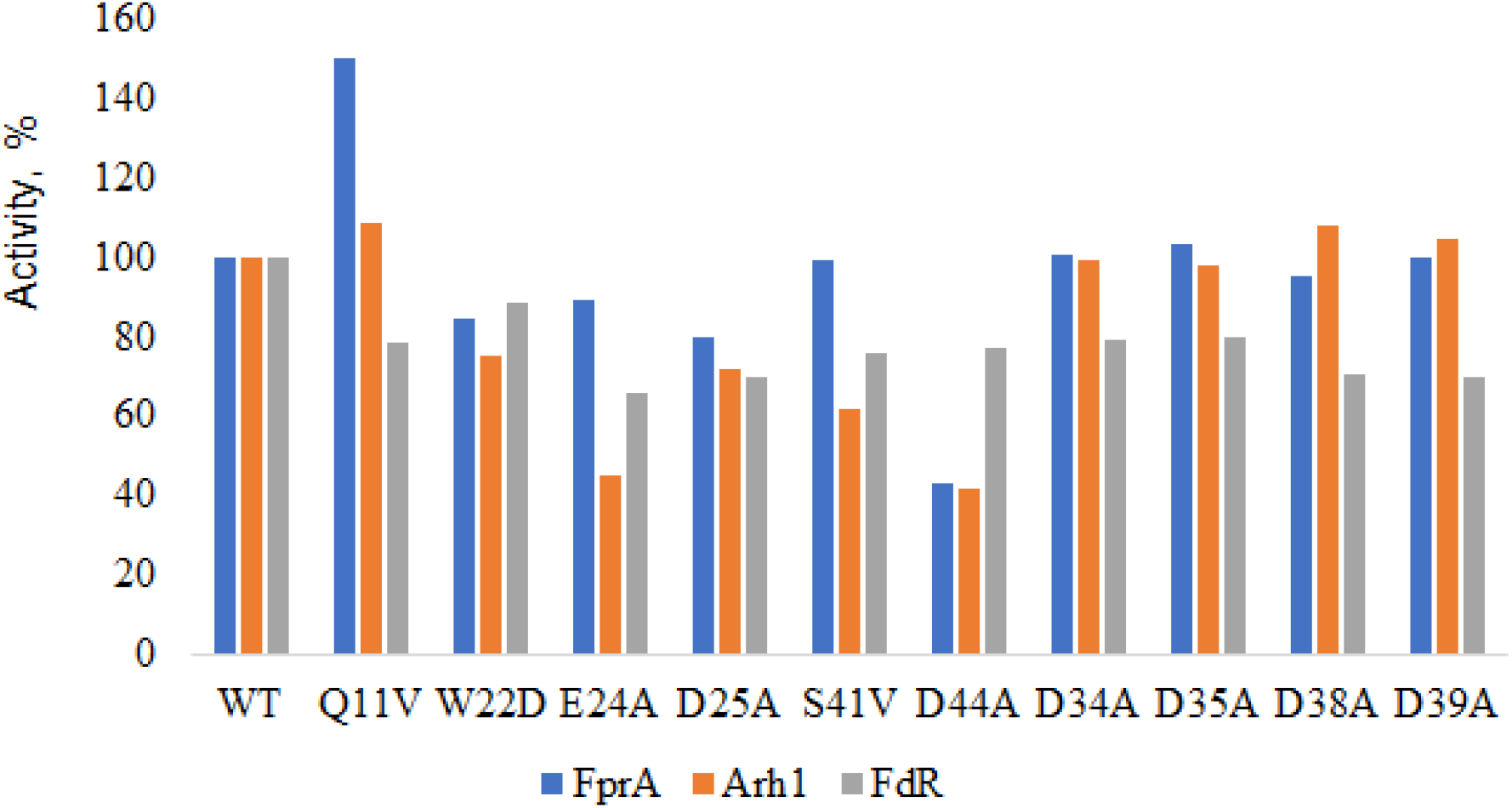
Catalytic activity of CYP124 reconstituted with RubB mutants in the system with Arh1 (blue bars), FprA (red bars), or FdR (grey bars).

We then compared the effects of RubB mutations on catalytic activity of CYP124, CYP125, and CYP142. We chose the cognate reductase FprA as a redox-partner, because catalytic activities of CYP125 and CYP142 was low in the reconstituted system containing FdR. The Q11V mutation, which introduces a residue conserved in other rubredoxins (Fig. S6), slightly increased catalytic activities of CYP124, CYP125, and CYP142. Similarly, D35 and D39 mutations led to an increase in CYP125 activity of about 1.5 fold but did not alter activities of CYP142 and CYP124 (Fig. 5). In the presence of the RubB D25A mutant, there was a 6.4-fold decrease in catalytic activity of CYP142 compared to activity in the presence of the wild type RubB, but CYP125 and CYP124 activities were unchanged (Fig. 5). The proximal surface of CYP142 is less positively charged than those of CYP124 and CYP125 (Fig. 6), suggesting that RubB D25A mutation affects the electrostatic complementarity between RubB and CYP142.

**Fig 5.**
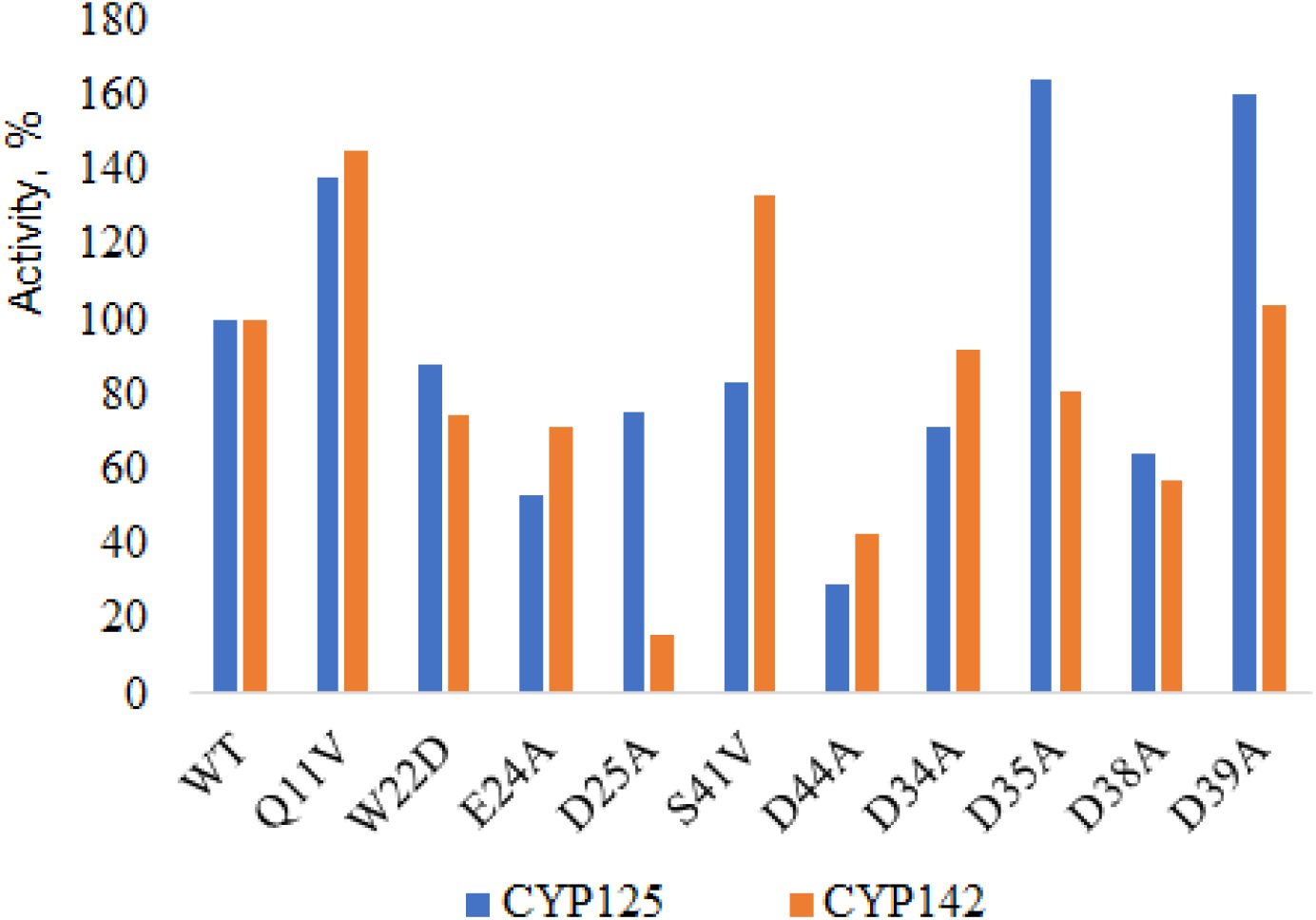
Catalytic activity of CYP125 (blue bars) and CYP142 (orange bars) reconstituted with FprA and indicated RubB mutants.

**Fig 6.**
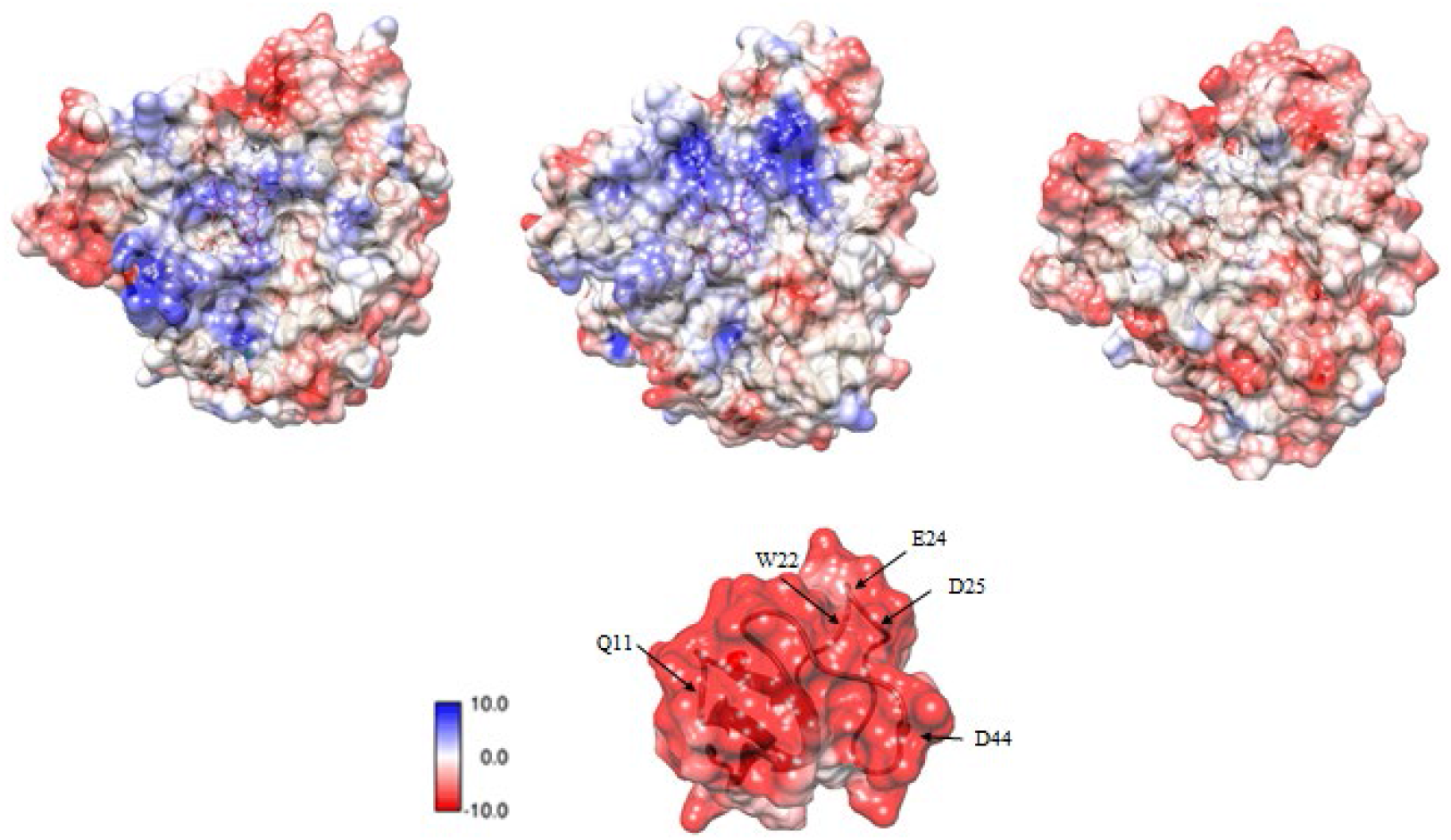
Electrostatic potential on the proximal surface of CYP124 (a), CYP125 (b), CYP142 (c) and RubB (d).

Mutations E24A, D44A, and D38A in RubB led to decreases in enzymatic activity of CYP125 and CYP142, with D44A the most influential reducing catalysis of all three CYPs (Fig. 5). Similar mutation of aspartate residue near iron-sulfur cluster was previously studied in the putidaredoxin from *P. putida* where D38A mutation resulted in almost complete loss of catalytic activity of CYP101A1 [27]. The effect of the D44A mutation was not as dramatic, suggesting rather low selectivity for RubB. These data support our hypothesis that D44 is involved in the interaction with reductase as sites of RubB interaction with both reductase and CYPs could be the same. Considering the small size of RubB it is reasonable to assume that it functions as a shuttle within CYP-dependent systems. RubB is very tolerant to amino acid substitutions as single charge neutralizations did not dramatically affect the CYP-RubB interactions. The RubB thus probably forms transient non-specific complexes within the CYP-dependent system.

## Discussion

Rubredoxins facilitate redox reactions and electron transfer as a single protein or as a domain within different protein families. Their functions vary from facilitation of adaptation to a changing redox environment [28] to the maintenance of overall protein stability [29] and are associated with different developmental processes [30]. In present study, we demonstrated that the Mtb rubredoxin RubB is a potent redox partner for Mtb CYPs. CYPs catalyze stereo- and regioselective oxidation of various endogenous substrates and produce bioactive compounds essential for medicine, agriculture and industry; they are indispensable components of biotechnological production of expensive or complex molecules [31]. The function of many CYPs of Mtb are unknown. The ability of RubB to transfer electrons from different reductases to at least three CYPs will make RubB an important research tool in the cytochrome P450 field.

Chemically modified electrodes such as those modified with carbon nanomaterials are used for the investigation of direct electrochemistry of metalloproteins. Composites based on carbon nanomaterials are promising for electrode modification to improve analytical parameters and sensor performance [32–34]. Based on our experimental data on electrochemistry of RubB carbon nanotubes are suitable promoters for negatively charged rubredoxin at carbon screen-printed electrodes. Carbon nanostructures possess electrochemically active surfaces and metallic or semiconductor properties that can induce catalysis by participating in electron transfer processes [35, 36]. An external supply of electrons using such electrodes via rubredoxin to CYPs may represent a new electron source for biomedical and biotechnological applications.

CYP124 catalyzes selective vitamin D3 hydroxylation, producing vitamin D3 metabolites with high biotechnological potential. An important focus of engineering of P450-dependent pathways is reducing equivalents to drive the catalytic cycle. The small size and high stability of RubB could be used in design of light-driven fusion proteins suitable for biotechnological applications, similarly to the fusion of ferredoxin and plant CYP79A1, which receives electrons directly from photosystem I [37]. Even more exciting from our point of view is the possibility of design of “infusion” CYPs. The rubredoxin fold retains electron transfer properties even in the context of an 18-amino acid peptide [38]. It may be possible to insert such a miniaturized rubredoxin into the meander region of CYP on the proximal surface (where the redox partner binds) to obtain infusion proteins suitable for a wide variety of applications.

Rubredoxin could also be used as a tag for screening of expression of colorless proteins [39], or to increase yields during expression of highly basic proteins.

Our finding that RubB is able to support catalytic activity of cholesterol-metabolizing CYPs furthers our understanding of these Mtb proteins. CYPs are essential for bacterial viability and pathogenicity [40]. A rubredoxin domain in the Mtb protein kinase G is important for kinase function under different redox conditions [28]. We suggest that rubredoxin motifs in Mtb proteins enable the bacterium to adapt to different microenvironments during its life cycle. In case of CYPs, RubB may participate in a switch mechanism from classic ferredoxins under iron starvation.

In summary, our studies revealed that RubB supports catalytic activity of heme-containing proteins of cytochrome P450 family in Mtb. From a broad perspective, our result have value in the biotechnology and synthetic biology applications of cytochrome P450 catalysis.

## Materials and methods

### Cloning and expression of Mtb RubB

The *RubB* (Rv3250c) gene was amplified from Mtb genomic DNA. The amplification reaction product was ligated into the expression vector pET11a.

*E. coli* C43 competent cells were transformed with pET11a containing *RubB*. Transformed cells were screened on Petri dishes with LB-agar containing ampicillin (100 μg/ml). An overnight culture (3 ml) was used to inoculate 0.5 L of TB-medium containing 100 mM potassium-phosphate buffer, pH 7.4, and 100 μg/ml ampicillin. The mixture was incubated in a thermostated orbital shaker (180 rpm) at 37°C. After absorbance at 600 nm reached ~ 0.4, RubB expression was induced by IPTG (final concentration 0.5 mM), additional FeCl3 (final concentration 100 μg/ml). After 20 h of incubation at 20°C with shaking at 100 rpm, the cells were collected by centrifugation (8000 g, 10 min). The pellet was resuspended in 50 mM potassium phosphate buffer, pH 7.4, containing 20% glycerol, 0.1 mM EDTA, and 0.5 mM PMSF. The cells were stored at –73 °C.

### Purification of RubB

The cell suspension was sonicated in an ice-water bath (7 x 1-min pulses with 1-min intervals). The suspension was centrifuged for 1 h at 20,500 rpm and the supernatant was applied to a DEAE-Sepharose column equilibrated with buffer A (50 mM potassium phosphate buffer, pH 7.4, containing 0.1 mM EDTA). The column was washed with 2-3 volumes of buffer A, and then with 10 volumes of buffer A containing 15 mM NaCl. RubB was eluted from the column with buffer A containing 200 mM NaCl. Eluted fractions were applied to a Superdex 75 16/60 column equilibrated with buffer A containing 200 mM NaCl. The colored band containing RubB was collected.

### Analytical methods

RubB purity was shown to be >95% by SDS–PAGE. Spectral studies were performed on a Cary200 spectrophotometer in 50 mM potassium phosphate buffer, pH 7.4. The concentration of RubB was determined using molar absorption coefficient of 6.9 mM^−1^ cm^−1^ at 490 nm [41].

### Electrochemical measurements

Electrochemical measurements were carried out using an EmStat3 potentiostat (PalmSens) equipped with the PSTrace software (version 5.4). Three-pronged electrodes obtained by screen-printing with graphite were used as working and auxiliary electrodes, and a silver chloride electrode was used as the reference electrode (http://www.colorel.ru). The diameter of the working electrode was 0.2 cm (area 0.0314 cm^2^). Electrochemical measurements were performed at room temperature in 0.1 M potassium phosphate buffer (pH 7.4), containing 0.05 M NaCl. Cyclic voltammograms were obtained at a scan rate of 0.05-0.1 V s ^-1^ and potential range from +0.5 to −0.8 V (vs. Ag/AgCl). All potentials are referred to relative to the Ag/AgCl reference electrode. The average values from three independent experiments are reported.

Prior to measurements, the SPEs were modified by aqueous dispersions of MWCNTs (Sigma-Aldrich) obtained from solubilized by amphiphilic poly(1,2-butadiene)-*block*-poly(2-(*N,N*-dimethylamino)ethyl methacrylate) (PB_290_-*b*-PDMAEMA_240_) diblock copolymer for a mean diameter of 9.5 nm and a length of 1 μm. A detailed description of the polymer synthesis, dispersion preparation, and the physico-chemical features of the modified SPEs can be found elsewhere [34], [42]. For preparation of the modified electrodes, 2 μL of the aqueous PB_290_-*b*-PDMAEMA_240_/MWCNT dispersion was dropped onto an area of the SPE and incubated for 15 min at 37 ºC until completely dry [33, 34].

For further incorporation of RubB, 1 μL of a solution of RubB at an appropriate concentration was applied to the surface of the modified SPE. The electrodes were allowed to stand for 12 h at 4 °C in a humid chamber. During the investigation of RubB potential, electrodes were in a planar regimen and covered by 60 μL of 0.1 M potassium phosphate buffer (pH 7.4), containing 0.05 M NaCl. Typical peak current separation (*ΔEp*) at 100 mV s^-1^ scan rate was 183 mV. The immobilized RubB showed two peaks, the reduction peak (the cathodic *E*pc) and the oxidation peak (the anodic *E*pa), in direct non-catalytic cyclic voltammetry with midpoint potentials from −231 mV to −264 mV. The midpoint potential *E*^0Ꞌ^ was calculated from the values of anodic and cathodic peak potentials using the equation *E*^0Ꞌ^ = (*E*_pc_ + *E*_pa_)/2.

### Circular dichroism spectroscopy

A calibrated Jasco Model J-820 spectropolarimeter was used to collect CD data from 0.3 mg/ml RubB samples in 50 mM potassium-phosphate buffer, pH 7.4. Spectra were recorded in a quartz cell of 1-cm path length between 200 and 260 nm at 298 K. Spectra were recorded in triplicate with a bandwidth of 1.0 nm and a time constant of 1.0 s, and the average was reported. The average spectra were processed by subtracting a blank spectrum from the protein spectrum.

### Isothermal titration calorimetry for RubB binding to CYP124

Experiments were performed using an ITC200 calorimeter equipped with the control and data acquisition and analysis software ORIGIN 7 (MicroCal Inc.). RubB and CYP124 were dialyzed overnight against 50 mM potassium-phosphate buffer, pH 7.4. An aliquot of 20 μM CYP124 was placed in the calorimetric cell and titrated with 1 mM RubB. The first injection (1 μl, omitted from the analysis) was followed by 20 injections of 4 μl at 2-min intervals. The titration syringe was continuously stirred at 750 rpm, and the temperature of the calorimetric cell was 25 °C. Injecting the RubB into the buffer alone was used as reference titration. When CYP124 substrate cholestenone was tested, cholestenone was dissolved in DMSO and was added to CYP124 at the final concentration 5 μM. DMSO was added to RubB in the same amount.

### Isothermal titration calorimetry for metal binding to RubB

RubB was dialyzed overnight against 50 mM HEPES, pH 7.4. RubB (0.2 mg/ml) was placed in the calorimetric cell and titrated with 200 mM zinc acetate dissolved in 50 mM HEPES, pH 7.4. The first injection (1 μl, omitted from the analysis) was followed by 20 injections of 4 μl at 2-min intervals. The titration syringe was continuously stirred at 750 rpm and the temperature of the calorimetric cell was 25°C. Injection of the zinc acetate into the buffer alone was used as a reference titration.

### Differential scanning calorimetry studies of RubB

DSC experiments were performed using a Microcal VP-DSC instrument (Malvern Instruments). The temperature gradient was 20 °C to 100 °C, the scan rate 60 °C/h. Baseline scans were of assay buffer (50 mM HEPES, pH 5.0). RubB was diluted to a final concentration of 50 μM in the same buffer. Zinc acetate was added to a final concentration of 10 mM. For data analysis Origin Software (OriginLab) was used.

### Catalytic activity of CYP124, CYP125, CYP142

Expression and purification of CYP124, CYP125, and CYP142 was performed as described previously [43]. Expression and purification of Arh1 and FprA was performed as described previously [44].

Catalytic activity of CYP124, CYP125 and CYP142 was determined in the reconstituted system containing 0.5 μM CYP, 1 μM FprA and 5 μM RubB in 25 mM HEPES, pH 7.4 containing 100 μM of 7-ketocholesterol (for CYP124) or cholestenone (for CYP142 and CYP125). Aliquots of concentrated proteins were mixed and pre-incubated for 5 min at room temperature. After incubation at 30°C for 5 min, the reaction was started by adding NADPH-regeneration system (3.3 mg/ml glucose-6-phosphate, 0.16 mg/ml NADPH, 0.6 μl/ml glucose-6-phosphate dehydrogenase). The mixture was incubated at 30 °C, and aliquots (0.5 ml) were taken at 15 and 30 min time intervals. Reactions were stopped by addition of 5 ml of methylene chloride, vortexed, and centrifuged at 3,000 rpm for 5 min. The organic layer was dried under argon flow, 100 μl of methanol was added to the pellet, and products were analyzed by HPLC using a C18 Luna 100 Å, 250 × 4.6 mm column on an Agilent Technologies 1200 Series instrument.

### Crystallization, data collection, X-ray crystallography, and modeling of Rub-CYP complex

Purified RubB was diluted to 1.5 mM in 10 mM Tris-HCl, pH 7.4 for crystallization trials. The initial screening was performed using commercially available screening kits from Qiagen and Molecular Dimensions in 96-well plate format using sitting-drop technique. Red-colored crystals of RubB grew in 2 days from 0.05 M zinc acetate, 20% PEG3350 at 20 °C. Optimization of crystallization conditions was carried out manually in 96-well plates using sitting-drop protocol. The best crystals are grown in 100 mM Tris-HCl pH 7.5, 50 mM zinc acetate, 25% PEG3350.

Diffraction data were collected at the European Synchrotron Radiation Facility (ESRF) beamline ID30B. The data collection strategy was optimized in BEST [45]. All data were processed in the XDS software package [46]. Processed data were corrected for anisotropy using the STARANISO server [http://staraniso.globalphasing.org/cgi-bin/staraniso.cgi]. A local mean *I*/*σ*(*I*) value of 1.20 was used to determine the anisotropic diffraction-limit surface.

The phase problem was solved in the automatic molecular replacement pipeline MoRDa [47]. The obtained space group was P1 with eight molecules per asymmetric unit. The model was rebuilt in phenix.autobuild [48]. Multiple rounds of model refinement with anisotropic B-factors were done in refmac5 [49] and phenix.refine [50]. Manual refinement was performed in *Coot* [51]. The quality of the resulting model was analyzed using phenix.molprobity [52] and QCCheck (https://smb.slac.stanford.edu/jcsg/QC/). The secondary structure was validated using the DSSP program [53]. Zinc ion geometry was determined using the CheckMyMetal web server [54, 55]. Data collection and final refinement statistics are presented in Table S1.

### Site-directed mutagenesis of RubB

Mutations were introduced by site-directed mutagenesis using oligonucleotides listed in Table S2. PCR conditions for single amino acid mutations were as follows: 25 cycles of 10 s at 98°C, 5 s at 60°C, followed by 6 min at 68°C. The resulting mutant plasmids were verified by DNA sequencing.

### Size-exclusion chromatography-multiple angle light scattering experiments

The experimental system consisted of a size-exclusion column and an HPLC equipped with a DAWN MALS detector (Wyatt Technology) and an Optilab differential refractive index detector (Wyatt Technology). RubB diluted in 50 mM HEPES, pH 7.4, containing 0.05 M NaCl to a final concentration of 1 mg/ml was applied to a Superdex 10/300 column (GE Healthcare) run at a flow rate of 0.5 ml/min. Zinc chloride was added to a final concentration of 5 mM if needed. Radius of gyration and absolute molecular mass were calculated using Wyatt’s ASTRA software.

## Abbreviations

CYP: cytochrome P450
RubB: rubredoxin B
Mtb: Mycobacterium tuberculosis
SPE: screen-printed electrode
MWCNT: multiwalled carbon nanotubes
FprA: NADPH-ferredoxin reductase
FdR - NADH: ferredoxin reductase
Arh1: adrenodoxin reductase-like flavoprotein
PETH: ferredoxin
NADP: reductase
ITC: isothermal calorimetry
CD: circular dichroism
DSC: differential scanning calorimetry
HPLC: high performance liquid chromatography
RMSD: root mean square deviation.

## Author contributions

Conceptualization, A.G., N.S.;

Methodology, V.B., V.S., K.T., A.G., N.S.;

Investigation, T.S., A.K., I.G., An.K., D.V., S.B., E.M., Al.K., R.M., L.S., N.S.;

Writing – Original Draft. T.S., D.V., N.S.;

Writing – Review & Editing, S.B., V.B., A.G., N.S.;

Funding Acquisition, V.B., A.G., N.S.;

Supervision, V.B., A.G., N.S.

## Acknowledgments

This study was inspired by personal communication with Prof. Trevor Forsyth (The Institut Laue– Langevin, France). This work was supported by a joint grant received from Belarusian Republican Foundation for Fundamental Research, B20R-061 and Russian Foundation for Basic Research, 20-54-00005. V.B. is supported by the Ministry of Science and Higher Education of the Russian Federation (agreement #075-00337-20-03, project FSMG-2020-0003). We thank Dr. Dmitry V. Pergushov and Dr. Felix H. Schacher for their help with electrode modifiers. The electrode modifiers were developed in the frames of Russian Science Foundation (RSF, project no. 18-44-04011) and the Deutsche Forschungsgemeinschaft (DFG, SCHA1640/18-1) within a joint RSF-DFG grant. We acknowledge the ESRF Structural Biology Group.

## Notes

**Conflict of interests**: The authors declare no conflict of interest.

### Competing Interest Statement

The authors have declared no competing interest.

