## Supplemental Information for "A new twist of rubredoxin function in *M. tuberculosis*"

**Fig. S1.** Sequence similarity network representation for rubredoxins from bacteria, archaea, and eukaryotes. Clusters are colored according to classes for Proteobacteria and according to phylum for others. Nodes corresponding to single sequences and two-node clusters are not shown. Sequences were obtained using iterative HMM search tool jackhmmer [https://www.ebi.ac.uk/Tools/hmmer/search/jackhmmer] with the Rv3250c sequence used as input. Only rubredoxin domain sequences were analyzed. Sequence were annotated using SeqScrub tool (http://www.gabefoley.com/seqscrub/) [56] and then used for sequence similarity network building with EFI-EST (https://efi.igb.illinois.edu/efi-est/). The E-value cutoff for the analysis was 10-23. Visualization was performed in Cytoscape.

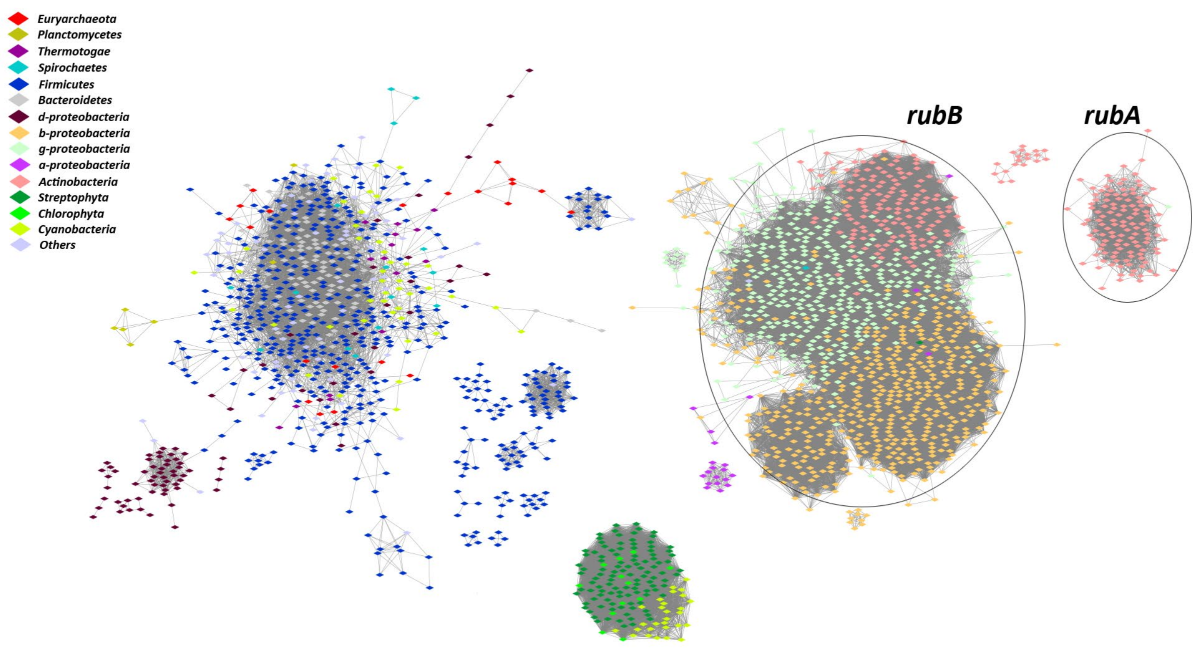

**Fig. S2.** Relative abundance of CYPs and redox partners across all bacteria. Sequences were obtained from the Proteomes database using jackhmmer tool. To evaluate the number of proteomes in each taxon, the numbers of dnaB, recA, and secA proteins were calculated and mean values were determined. The content of the proteins of interest was then compared to the mean number of proteomes. Fdx column includes sequences with single Fer4 (PF00037, two Fe-S clusters) or Fer4_15 (PF13459, one Fe-S cluster) domain corresponding to the two types of Mtb ferredoxins. Flavodoxins are represented as two separate columns, Fld and FMN_red. The Fld column includes proteins with any of five annotated flavodoxin domains Flavodoxin_1 (PF00258), Flavodoxin_2 (PF02525), Flavodoxin_3 (PF12641), Flavodoxin_4 (PF12682), or Flavodoxin_5 (PF12724). The FMN_red column includes homologs of Rv2771c of Mtb, which is a predicted flavodoxin (PF03358). The Rub column includes sequences with annotated rubredoxin domains (PF00301). Heatmap visualization was constructed using numpy, matplotlib, and seaborn libraries in python3.

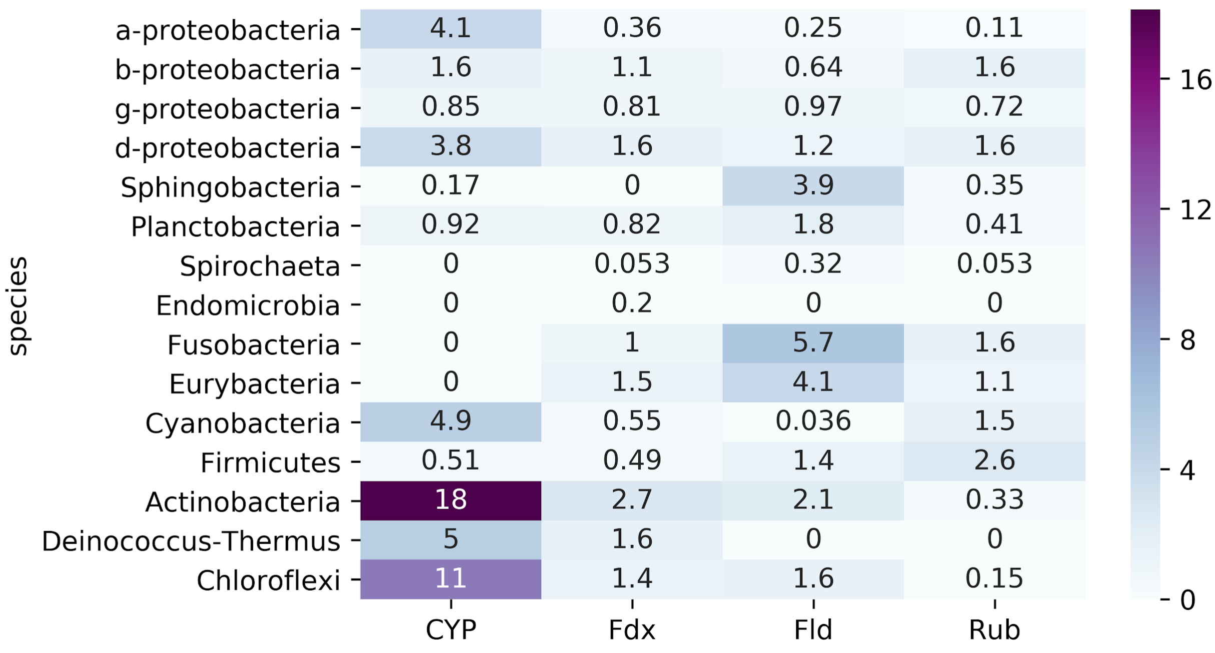

**Fig. S3.** CD spectrum of purified RubB. CD spectrum was recorded in 50 mM potassium-phosphate buffer, pH 7.4.

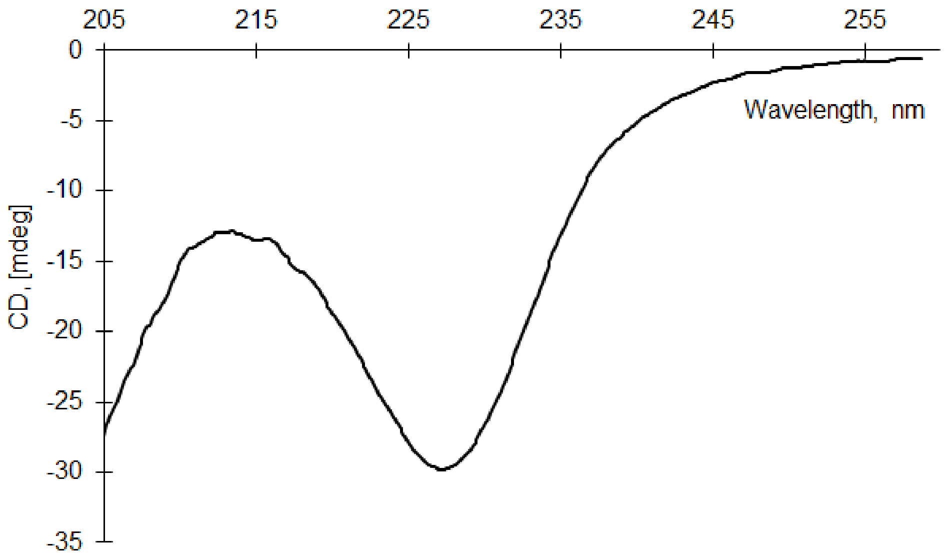

**Fig. S4.** Crystal packing of RubB. There are eight protein molecules in the asymmetric unit: A (yellow), B (red), C (cyan), D (blue), E (orange), F (green), G (grey-green), and H (magenta). Iron and zinc ions are depicted as orange and grey spheres, respectively.

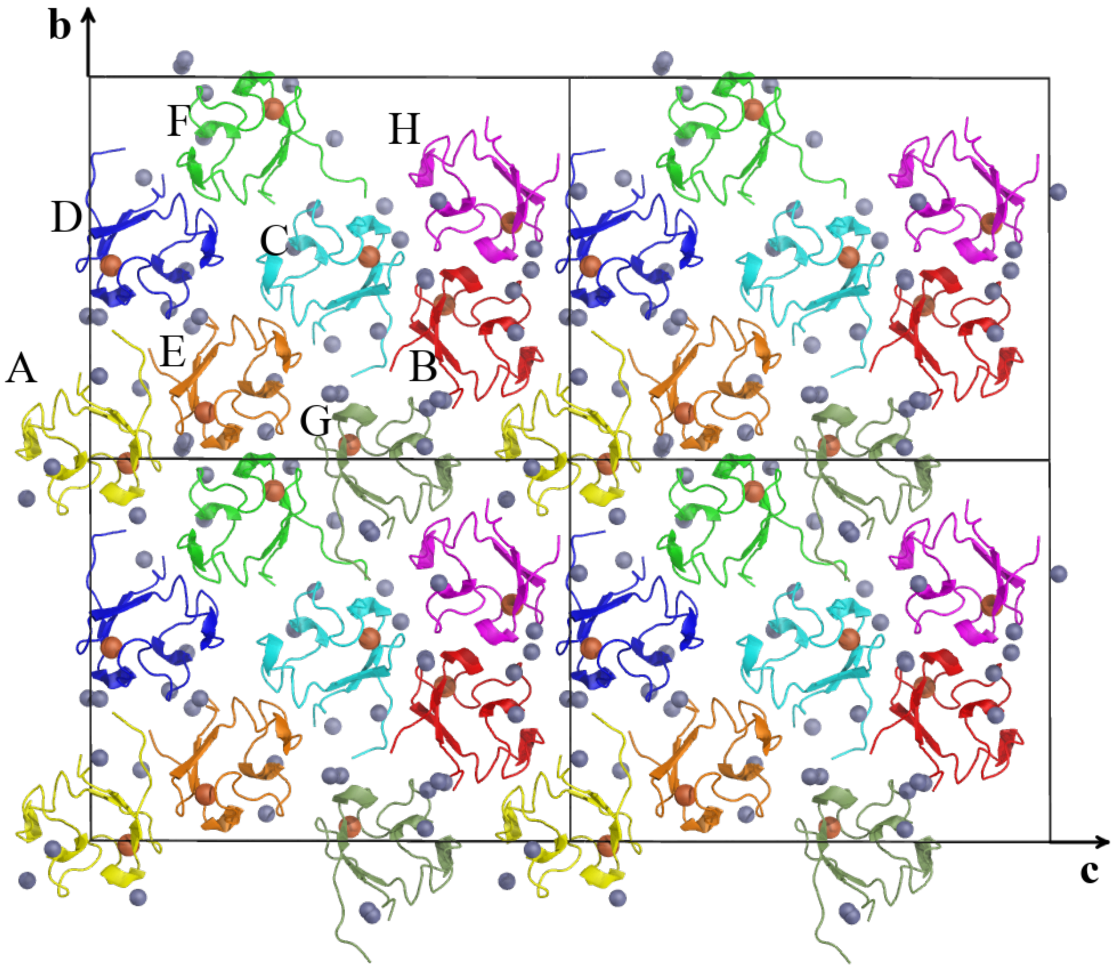

**Fig. S5.** ITC binding curve of RubB with $Zn^{2+}$ (injectant) in 50 mM HEPES, pH 7.4 at 25°C.

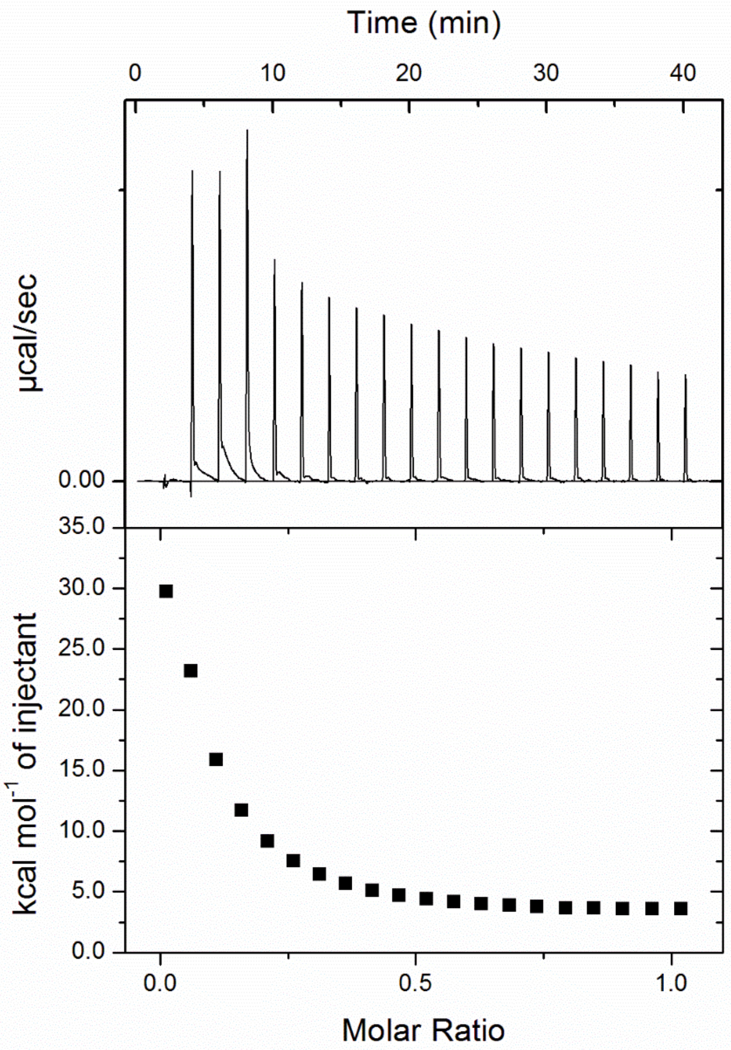

**Fig. S6.** Multiple sequence alignment of rubredoxins from *Clostridium cellulolyticum (Cce), Clostridium sticklandii (Cst), Clostridium pasteurianum (Cpa), Clostridium perfringens (Cpe), Clostridium thermosaccharolyticum (Cth), Megasphaera elsdenii (Me), Peptostreptococcus asaccharolyticus (Pas), Butyribacterium methylotrophicum (Bm), Heliobacillus mobilis (Hm), Chlorobium limicola (Chl), Desulfovibrio desulfuricans (Dd), Desulfovibrio vulgaris Hildenborough (DvH), Desulfovibrio vulgaris Miyazaki (DvM), Pyrococcus furiosus (Pf), Pyrococcus abbyssi (Pa), Desulfovibrio gigas (Dg), Pseudomonas aeruginosa (Pae), MtbB* - rubredoxin B, and *MtbA* - rubredoxin A. Conserved cysteines in CXXC motifs are colored yellow. Amino acids that form the hydrophobic core are colored green. Amino acids chosen for mutations based on multiple sequence alignment are colored red. Multiple sequence alignment was performed using ClustalW [57].

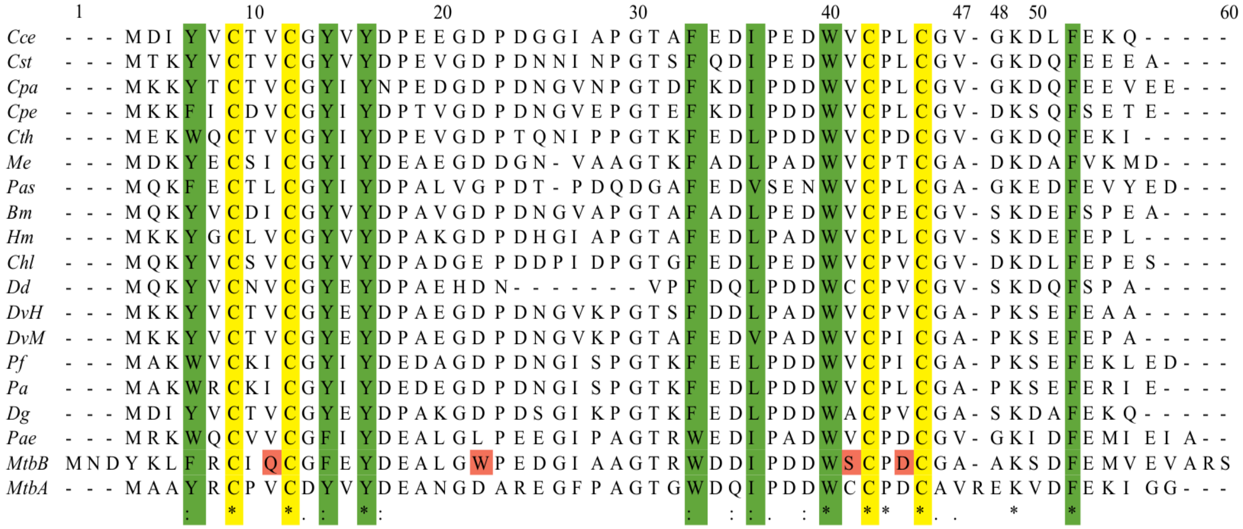

**Fig. S7.** Comparison of the active sites in X-ray and NMR structures of RubB. D44 in NMR models (colored green) have different side-chain rotamers than D44 from the X-ray structure (colored blue). Cα superimposition is based on residues C42-C45.

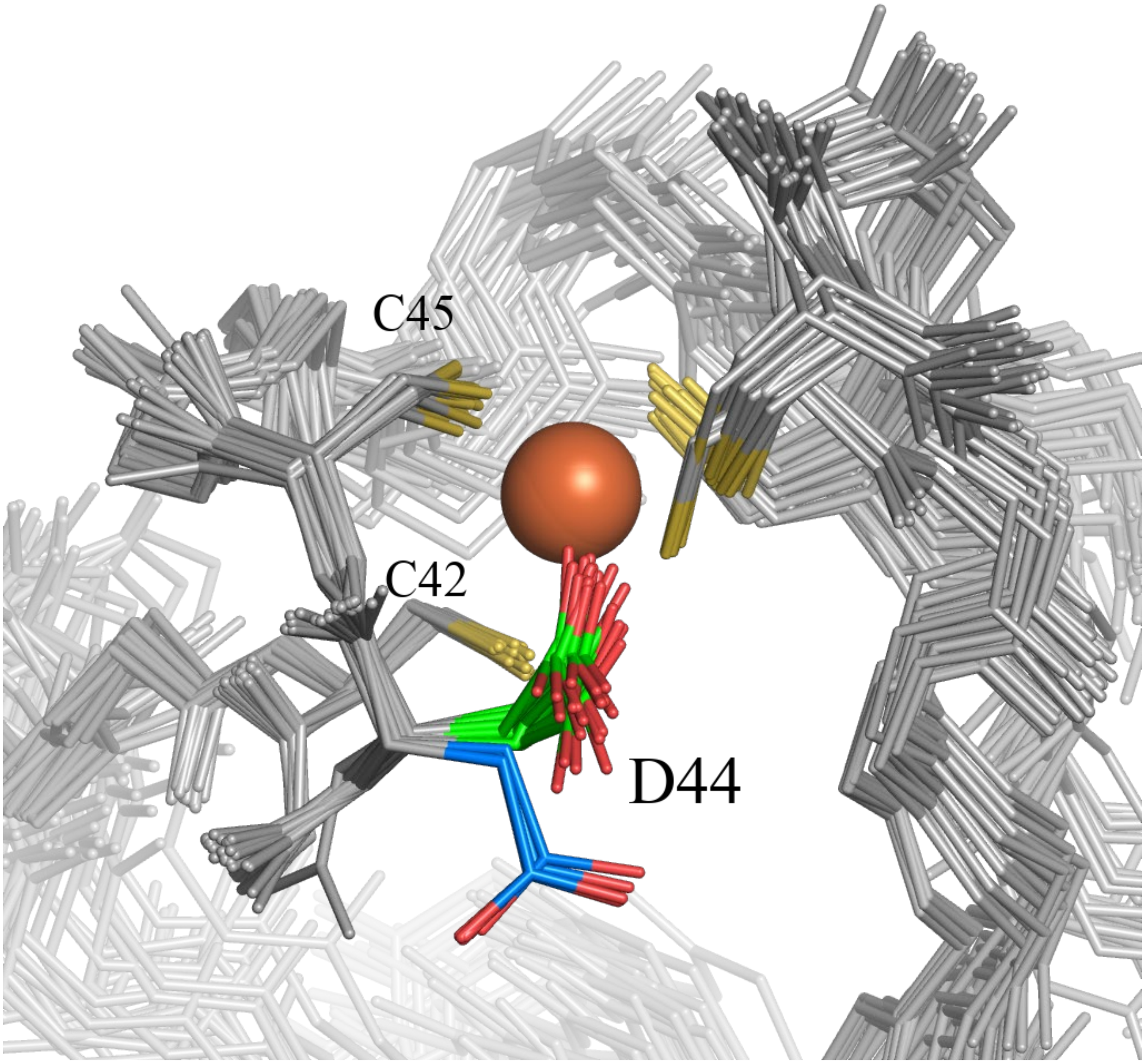

**Fig. S8.** DSC thermograms for RubB and mutated proteins.

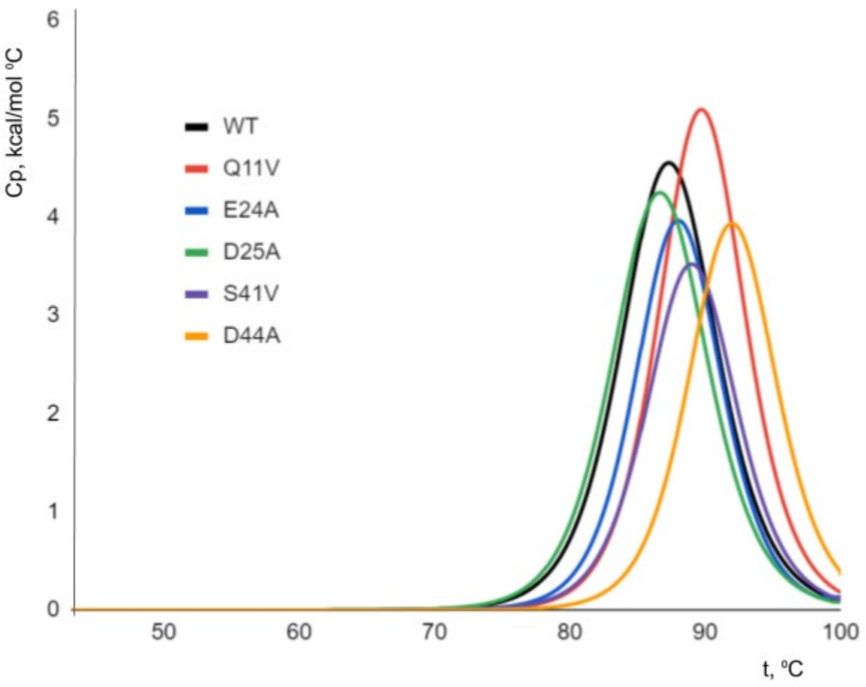

**Table S1.** Data collection and refinement statistics for RubB.

| RubB | | |
|---|---|---|
| **Data collection** | | |
| | Aimless | Staraniso |
| Wavelength, Å | 0.97625 | |
| Space group | P 1 | |
| Unit cell | 21.0 61.3 76.9 90.2 90.3 93.8 | |
| Resolution range, Å | 47.95 - 1.17 (1.31 - 1.17)[a] | |
| Resolution limits, Å | 1.17 | 1.15; 1.33; 1.57 |
| Total reflections | 411173(114440) | 263480(13312) |
| Unique reflections | 119411(33126) | 76442(3823) |
| Multiplicity | 3.5(3.5) | 3.4(3.5) |
| Completeness (%) | 93.18(90.95) | 59.7(10.5) |
| ellipsoidal (%) | | 88.4(60.2) |
| Mean I/sigma(I) | 4.69(0.55) | 7.0(1.5) |
| R-merge | 0.135(2.077) | 0.101(0.787) |
| R-meas | 0.160(2.466) | 0.121(0.933) |
| R-pim | 0.086(1.312) | 0.064(0.495) |
| CC1/2 | 0.994 (0.295) | 0.994(0.565) |
| **Refinement** | | |
| Resolution range, Å | 47.95 - 1.17 | |
| Reflections used in refinement | 76432 | |
| Reflections used for R-free | 3786 | |
| R-work | 0.154 | |
| R-free | 0.187 | |
| Number of non-hydrogen atoms | 4742 | |
| macromolecules | 4002 | |
| ligands | 116 | |
| solvent | 624 | |

| | |
|---|---|
| Protein residues | 471 |
| RMS(bonds) | 0.004 |
| RMS(angles) | 0.66 |
| Ramachandran favored (%) | 98.46 |
| Ramachandran outliers (%) | 0.00 |
| Rotamer outliers (%) | 0.46 |
| Clashscore | 1.39 |
| Average B-factor | 13.47 |
| macromolecules | 11.46 |
| ligands | 29.79 |
| solvent | 23.34 |

[a]Statistics for the highest resolution shell are shown in parentheses.

**Table S2.** Primers used to obtain RubB mutants.

| Mutation | Primers sequence |
|---|---|
| D44A | Fwd: 5'-CTGTTCCGCTGTATCGTATGCGGCTTTGAGTAC-3'<br>Rev: 5'-GTACTCAAAGCCGCATACGATACAGCGGAACAG-3' |
| Q11V | Fwd: 5'-CTGTTCCGCTGTATCGTATGCGGCTTTGAGTAC-3'<br>Rev: 5'-GTACTCAAAGCCGCATACGATACAGCGGAACAG-3' |
| S41V | Fwd: 5'-CCCCGATGACTGGGTTTGCCCGGATTG-3'<br>Rev: 5'-CAATCCGGGCAAACCCAGTCATCGGGG-3' |
| E24A | Fwd: 5'-CGATGCCGTCCGCCGGCCAACCC-3'<br>Rev: 5'-GGGTTGGCCGGCGGACGGCATCG-3' |
| D25A | Fwd: 5'-CCGCGATGCCGGCCTCCGGCCAA-3'<br>Rev: 5'-TTGGCCGGAGGCCGGCATCGCGG-3' |
